# Improvement of Spontaneous Locomotor Activity in a Murine Model of Duchenne Muscular Dystrophy by N-Acetylglucosamine Alone and in Combination with Prednisolone

**DOI:** 10.1101/2024.08.25.609562

**Authors:** Masahiko. S. Satoh, Guillaume St-Pierre, Ann Rancourt, Maude Fillion, Sachiko Sato

## Abstract

N-acetylglucosamine (GlcNAc) is an endogenous compound whose intracellular concentration is closely associated with the biosynthesis of acetyllactosamine-rich N-linked oligosaccharides. These oligosaccharides interact with mammalian lectin galectin-3, mediating cell surface receptor dynamics as well as cell-to-cell and cell-to-extracellular matrix interactions. Our previous and recent studies suggest that GlcNAc, in conjunction with galectin-3, augments muscle regeneration *in vitro*. We have also demonstrated that intraperitoneal GlcNAc administration improves muscle strength in a murine model of Duchenne muscular dystrophy (DMD) (*mdx* mice). Here, we show that oral administration of GlcNAc significantly improves the spontaneous locomotor activity of mdx mice. Administering GlcNAc at concentrations of 0.6, 1.2, 1.8, and 2.4 g/kg body weight per day for 35 days significantly improved nocturnal spontaneous locomotor activity at all those doses, with the 1.2 g/kg body weight dose reducing damages of extensor digitorum longus muscle by nearly 50%. While consecutive forced exercises, including horizontal and downhill treadmill running, reduced GlcNAc-promoted locomotor activity, treatment with 0.6 and 1.2 g/kg body weight treatment results in increased spontaneous locomotor activity. These results suggest that GlcNAc enhances overall muscle health, likely through promoting muscle repair/regeneration rather than preventing damage formation. Notably, co-administration of GlcNAc with prednisolone, a corticosteroid commonly used in DMD patients, further enhanced spontaneous locomotor improvement in *mdx* mice compared to prednisolone alone. These findings suggest that GlcNAc has the potential to improve the clinical status of DMD patients, either as a monotherapy or in combination with corticosteroids.

## Introduction

Duchenne muscular dystrophy (DMD) is an inherited X-linked disorder characterized by progressive muscle wasting, affecting approximately 1 in 3,500 male births worldwide(1–3). DMD is caused by mutations in the dystrophin gene, which encodes the dystrophin protein, a crucial component of the dystrophin-associated glycoprotein complex (DAG) that connects the actin cytoskeleton to the extracellular matrix (3–7). The absence or malfunction of dystrophin leads to the reduction of surface DAG, thereby compromising the integrity of muscle fibers, making them susceptible to damage and leading to chronic inflammation, fibrosis, and ultimately, muscle degeneration (1, 3, 5–9).

Corticosteroids, such as prednisolone, are commonly administered to DMD patients to reduce inflammation and slow disease progression, though they are associated with significant side effects (10–12). Recent therapeutic approaches include gene therapy, exon skipping, and the development of novel corticosteroids like vamorolone and deflazacort, which aim to minimize side effects while preserving anti-inflammatory efficacy (10–12) Previously, we reported that intraperitoneal administration of N-acetylglucosamine (GlcNAc) improved the muscular strength of mdx mice, a well-established mouse model of Duchenne muscular dystrophy (DMD) (13). This finding suggests that GlcNAc may have potential as a therapeutic agent for DMD patients, either as a standalone treatment or in combination with other DMD interventions Given the necessity of daily administration, oral delivery of GlcNAc is particularly advantageous for children with DMD. However, it was previously unknown whether oral administration could effectively mitigate disease progression. In this study, we investigate the therapeutic potential of oral GlcNAc as a standalone treatment and in combination with prednisolone in mdx mice.

GlcNAc is an endogenous compound that serves as a precursor for UDP-GlcNAc, which acts as a substrate for various N-acetylglucosaminyltransferases involved in protein glycosylation. These oligosaccharide-processing enzymes, including mannosyl glycoprotein N-acetylglucosaminyltransferases (MGATs), play a crucial role in the biosynthesis of N-linked oligosaccharides within the secretory pathway (14, 15). The increase in intracellular GlcNAc and UDP-GlcNAc, which can be augmented by GlcNAc administration, promotes the production of glycoproteins that carry acetyllactosamine-rich oligosaccharides (14, 15). These oligosaccharides play a critical role in the regulation of cell-cell and cell-cell matrix interaction as well as the dynamics of various membrane glycoproteins; this is partly mediated by galectin-3 (Gal-3), which specifically binds to those oligosaccharides (16–24). Our previous and recent studies suggest that interaction between oligosaccharide and galectin-3 promotes myogenesis (13, 25). Given that differentiating myoblasts and skeletal muscles of mdx mice and DMD patients express high levels of Gal-3(26–28), the interaction between the oligosaccharides and Gal-3 is likely important in myogenesis.

Orally administered GlcNAc is rapidly absorbed in the upper gastrointestinal tract and enters the circulation as GlcNAc (29–31). Previous preclinical toxicity studies in rats demonstrated that chronic oral administration of GlcNAc 2.5 g/kg body weight (BW) per day (equivalent to 0.6 g/kg BW per day in humans) for 52 weeks results in no adverse effects or histopathological changes on tissues (32–34). Additionally, GlcNAc demonstrated safety in patients with inflammatory bowel disease and multiple sclerosis at doses of 6 g and 12 g per day for 4 weeks with no reported adverse effects (35, 36). GlcNAc is a water-soluble monosaccharide with a light, sweet taste, likely contributing to high patient acceptance of its oral administration if therapeutic efficacies are demonstrated.

Here we report that oral administration of GlcNAc for 35 days can improve the spontaneous locomotor activity of *mdx* mice. Oral GlcNAc administration of 0.6, 1.2, 1.8, and 2.4 g/kg BW (equivalent to doses of 72, 144, 216, and 288 mg/kg BW per day, respectively, in humans (34)), did not affect BW or muscle mass in *mdx* mice. Spontaneous locomotor activity in *mdx* mice improved significantly at all doses of GlcNAc, and muscle damage was reduced with the administration of 1.2 g/kg BW GlcNAc, while creatine phosphokinase levels remained unchanged. To determine whether GlcNAc treatment preserves muscle health against forced exercises, mice were subject to consecutive forced exercises including horizontal and downhill treadmill running, during the final week of the 35-day treatment. Although those exercises reduced GlcNAc-promoted locomotor activity, treatment with 0.6 and 1.2 g/kg BW still increased locomotor activity even after forced exercises. Moreover, co-administration of GlcNAc with prednisolone further enhanced nocturnal locomotor activity. These findings suggest that while the optimal dose of GlcNAc for DMD patients remains to be determined, GlcNAc could be a promising therapeutic agent for improving muscle function in DMD, either as a standalone treatment or in combination with other interventions.

## Results

### Effect of GlcNAc on BW, Muscle Mass, and Creatine Phosphokinase Release (Protocol 1)

To evaluate the effects of GlcNAc on *mdx* mice, mice received GlcNAc continuously through their drinking water at the concentrations of 2.4, 4.8, and 9.6 mg/ml, which is equivalent to 0.6, 1.2, and 2.4 g/kg BW per day, respectively. Over the course of 35 days, no significant differences were observed in BW (Fig. 1A) or mortality (Fig. 1B), indicating that GlcNAc administration does not adversely affect the growth or survival of *mdx* mice. After 35 days, the masses of the Tibialis Anterior (TA), Extensor Digitorum Longus (EDL), and Soleus (Sol) muscles were measured. No significant differences were found in the absolute mass of the TA (Fig. 1C) or in the TA mass relative to BW (Fig. 1D) between treatment groups. Similarly, no significant changes were observed in the absolute or relative masses of the EDL (Fig. 1E and 1F) and Sol (Fig. 1G and 1H) muscles. These data suggest that GlcNAc does not influence muscle quantity. Additionally, while muscle damage in *mdx* mice is indirectly correlated with elevated plasma levels of creatine phosphokinase(37, 38), GlcNAc treatment did not alter these levels (Fig. 1I), suggesting that GlcNAc does not prevent the formation of muscle damage.

**Fig. 1.**
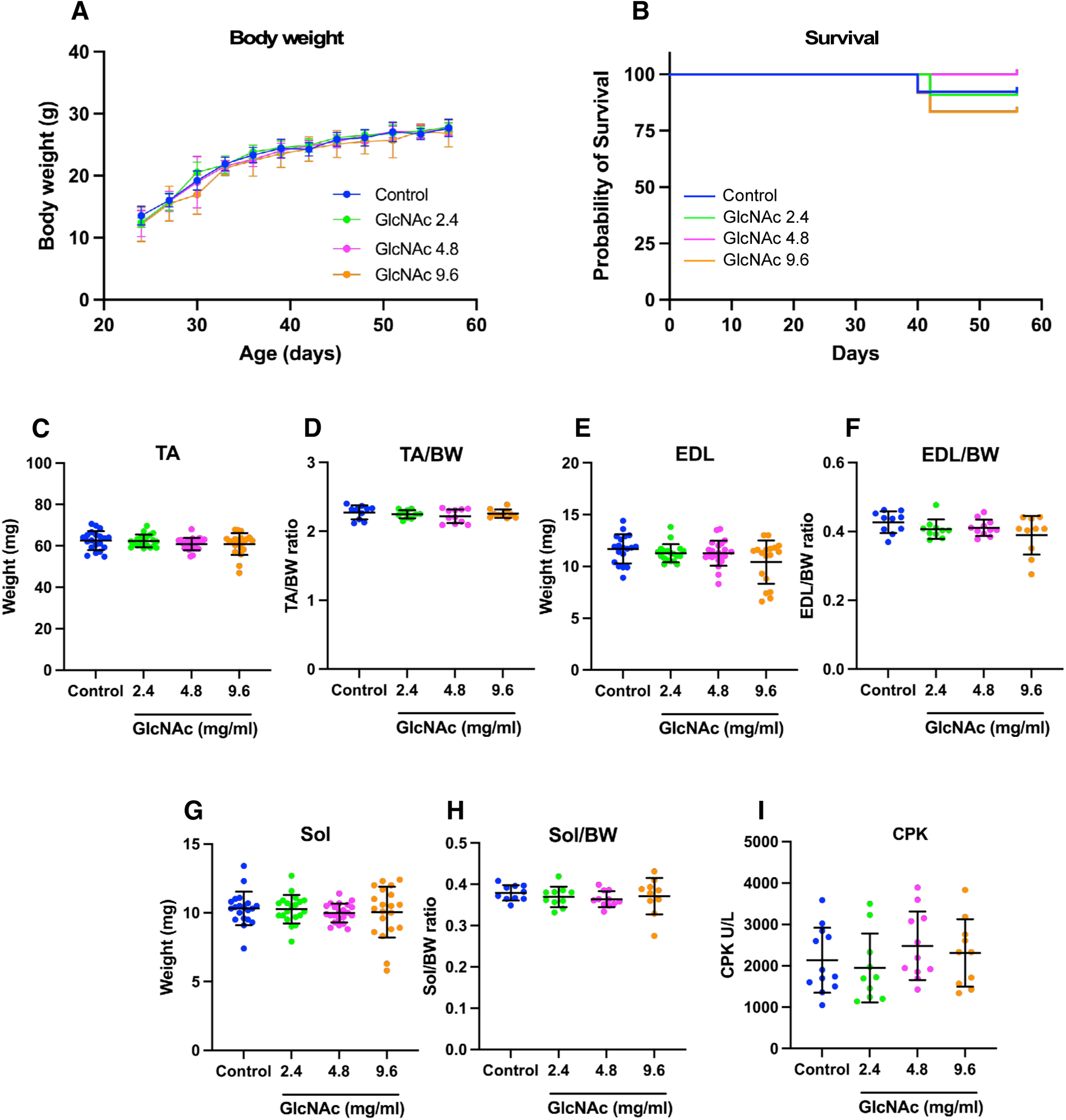
Impact of GlcNAc on BW, survival, and muscle mass in *mdx* mice (Protocol 1) GlcNAc (2.4, 4.8, and 9.6 mg/ml) was administered orally via voluntary intake to *mdx* mice. Mice were housed for 35 days, during which their body weight (BW) was regularly measured. The survival rate of the mice was monitored throughout the study. For both **A** and **B**, the number of mice used was 10. **C-H.** On day 35, the *mdx* mice were sacrificed, and the mass of the tibialis anterior (TA) (**C**), TA mass relative to BW (**D**), extensor digitorum longus (EDL) mass (**E**), EDL mass relative to BW (**F**), soleus (Sol) mass (**G**), and Sol mass relative to BW (**H**) were measured. For **C, E,** and **G**, the sample size was 20, and for **D, F,** and **H,** the sample size was 10. **I.** Creatine phosphokinase (CPK) levels in the serum were also measured (n = 10). Statistical analyses were performed using ordinary one-way ANOVA with Tukey’s post-hoc test (**A**, and **C-I)**, and Mantel-Cox test (**B**). No significant differences were observed. **A**, and **C-I**. Data represent means ± standard deviations.

### Histological Analysis of EDL

Next, we analyzed the muscle fiber size of EDL using Hematoxylin-Eosin (H&E) stained sections. To objectively assess muscle fiber size, we employed our in-house software, which converts H&E-stained images into grayscale, performs image segmentation to identify muscle fibers, and quantifies the areas of these fibers (Fig. 2A). Based on their size, muscle fibers were color-coded, as illustrated in Fig. 2A. Using this software, we investigated the effect of GlcNAc on muscle fibers but found no significant differences (Fig. 2B). We then quantitatively evaluated the formation of matrix space and inflammatory areas using the same software, which generates grayscale images and performs image segmentation to identify damaged areas (Fig. 2C). Combined analysis of the matrix space and inflammatory sites revealed significantly less (∼50%) damage in the EDL muscles of *mdx* mice treated with 4.8 mg/ml GlcNAc (Fig. 2D). The area of inflammation was also reduced (Fig. 2E), although no significant reduction was observed in the matrix space (Fig. 2F). This reduction in muscle damage occurred despite the lack of significant changes in creatine kinase levels (Fig. 1I). Given that our previous studies indicate that GlcNAc augments myogenesis *in vitro* (13, 25), these results suggest that GlcNAc may accelerate muscle regeneration rather than prevent the initial formation of muscle damage.

**Fig. 2.**
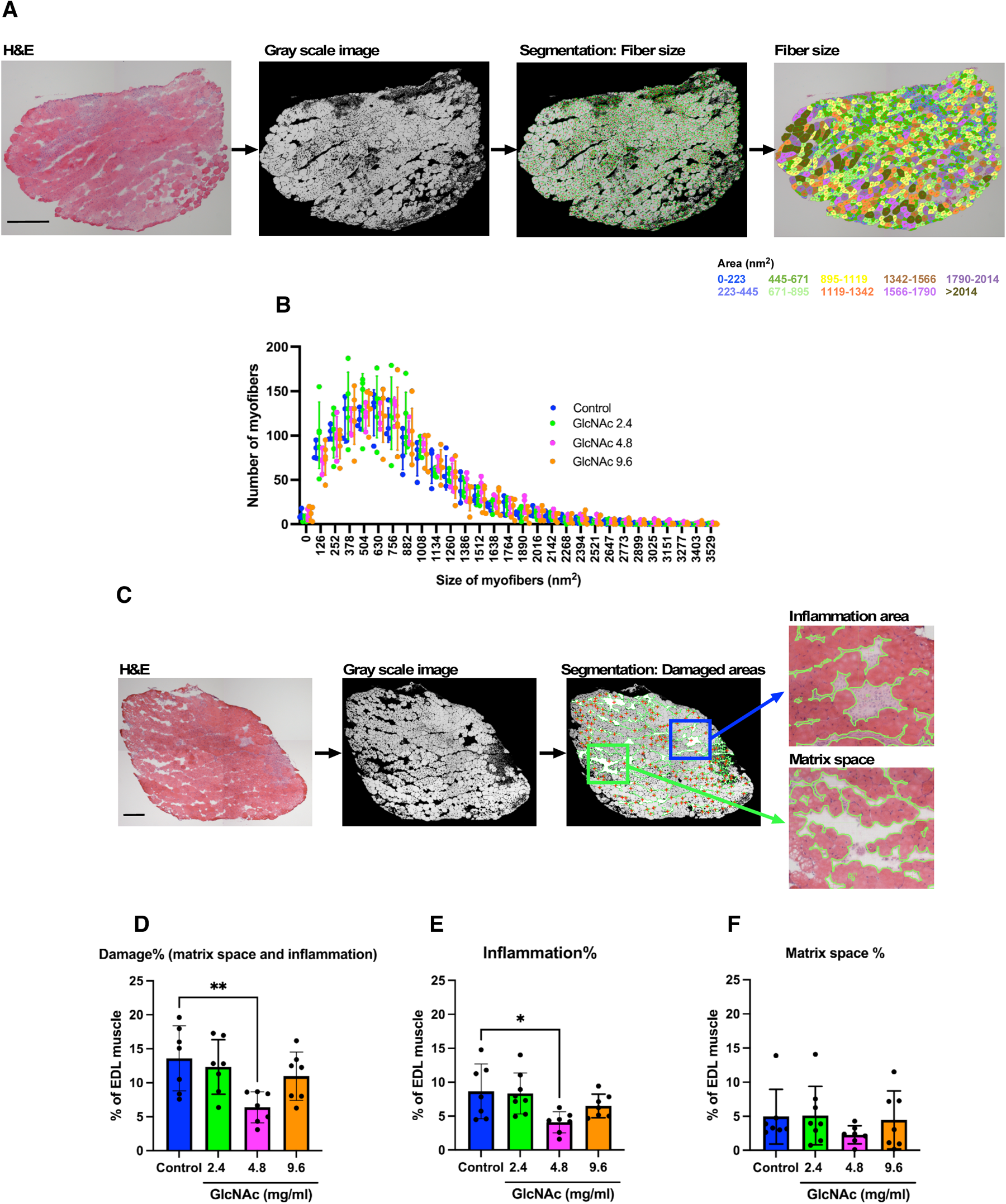
Effect of GlcNAc on muscular damage formation (Protocol 1) **A.** Tissue sections of the extensor digitorum longus (EDL) muscle were prepared and stained with hematoxylin and eosin (H&E). Images were captured and converted to grayscale for segmentation analysis using in-house software to measure muscle fiber sizes, which were illustrated using a color-coding system. **B.** The effect of GlcNAc on the muscle fiber size of EDL was analyzed. Statistical analysis was performed using two-way ANOVA, and no significant differences were observed. Data represent means ± standard deviations, with a sample size of 4 or 5 for each group. **C.** Images of the EDL sections stained with H&E were captured and converted to grayscale for segmentation analysis using in-house software to measure the areas of inflammation and matrix space. The detected areas used for quantification are indicated by green lines. **D-F.** The percentage of damaged area was calculated by combining the inflammation area and matrix space, normalized to the total area of the EDL (**D**). The percentage of the inflammation area (**E**) and matrix space (**F**) was calculated by normalizing these areas to the total area of the EDL. Statistical analysis was performed using one-way ANOVA with Dunnett’s test. Significance levels are indicated as *P < 0.01, and **P < 0.005. Data represent means ± standard deviations, with a sample size of 7 for each group. **A** and **C.** Scale bars indicate 100 nm.

### Treadmill Exercise in *mdx* Mice (Protocol 2)

To further assess whether GlcNAc influences muscle damage formation and test additional GlcNAc dose, we implemented Protocol 2 in which *mdx* mice were treated for 35 days with GlcNAc at concentrations of 2.4, 4.8, 7.2, and 9.6 mg/ml and subjected to treadmill running. Three weeks after the initiation of GlcNAc administration, mice treadmill session began in the 4th week (Fig. 3A). To acclimate the mice to treadmill running, training sessions were conducted on a flat treadmill at a speed of 5 m/min for 10 minutes on Day 1. After a day of rest, the speed was increased to 10 m/min for 10 minutes on Day 3. Following another day of rest, the speed was further increased to 15 m/min on Day 5. After two more days of rest, the final treadmill run was performed on Day 8, with a gradual increase in speed from 8 to 15 m/min over 49 minutes on a 15-degree downhill incline. After the run, Evans Blue Dye (EBD) was injected, and the mice were sacrificed the following day. No significant differences in BW (Fig. 3B) or survival (Fig. 3C) were observed between groups, and no significant differences were observed in the TA (Fig. 3D and E), EDL (Fig. 3F and G) or Sol (Fig. 3H and I) muscle masses, whether absolute or relative to BW. Additionally, the percentage of completed treadmill exercises did not differ significantly between groups (Fig. 3J). These results suggest that, under the conditions used, the lack of dystrophin did not prevent some *mdx* mice from completing downhill exercise. Furthermore, there were no significant differences between GlcNAc-treated and control mice in their ability to perform forced running.

**Fig. 3.**
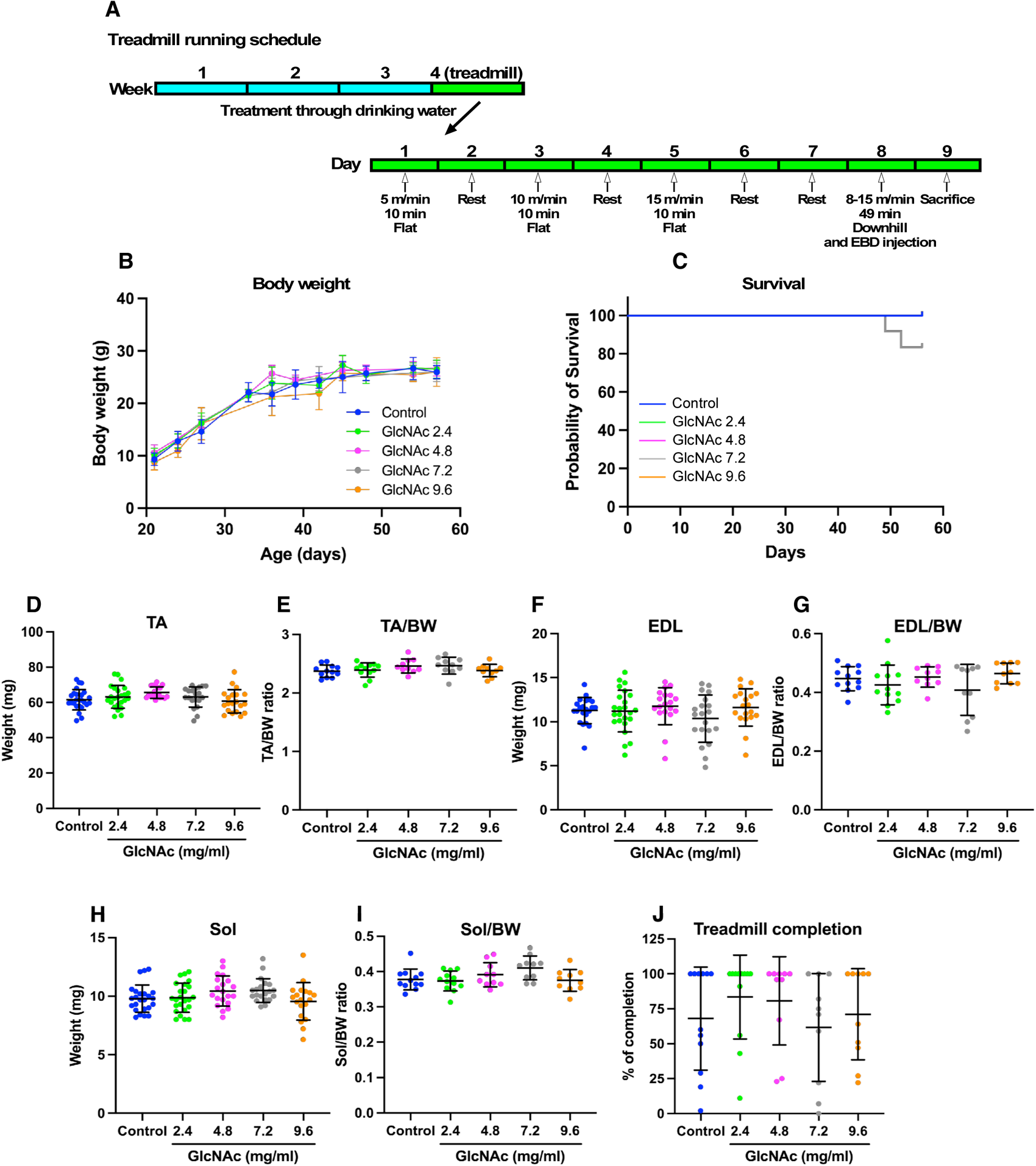
Impact of GlcNAc on BW, Survival, and Muscle Mass in *mdx* Mice Subjected to Treadmill Running (Protocol 2) **A.** Treadmill exercise schedule. After 26 ± 1 days of treatment, *mdx* mice were acclimated to treadmill running with three sessions on a flat treadmill, as illustrated. On Day 8, the mice were subjected to a faster treadmill run on a 15-degree downhill incline, with speeds gradually increasing from 8 to 15 m/min over 49 minutes. Immediately after the run, Evans Blue Dye (EBD) was injected to stain damaged muscles, and the mice were sacrificed the following day. Mice’s body weight (BW) was regularly measured throughout the study. **C.** The survival rate of the mice was monitored. For both panels **B** and **C**, the number of mice in the Control, 2.4, 4.8, 7.2, and 9.6 mg/ml GlcNAc-treated groups were 13, 14, 10, 11, and 10, respectively. **D-I.** On the day following the treadmill run (Day 8), mice were sacrificed, and the mass of the tibialis anterior (TA) (**D**), TA mass relative to BW (**E**), extensor digitorum longus (EDL) mass (**F**), EDL mass relative to BW (**G**), soleus (Sol) mass (**H**), and Sol mass relative to BW (**I**) were measured. For panels **D**, **F**, and **H**, the sample size was 20 to 24, and for panels **E**, **G**, and **I**, the sample size was 10 to 12. **J.** The percentage of mice that completed the treadmill run performed on Day 8 was calculated (sample size: 10 to 12). Statistical analyses were performed using ordinary one-way ANOVA with Tukey’s post-hoc test (**B**, and **D-J**) and the Mantel-Cox test (**C**). No significant differences were observed. Data in panels **B** and **D-J** represent means ± standard deviations.

### Muscle Damage Induced by Treadmill Running (Protocol 2)

Quantitative analysis of damage sites using H&E-stained sections of the EDL muscle following treadmill running revealed that muscle damage occurred across all GlcNAc-treated groups, with no significant differences from non-treated mice (Fig. 4A, B, and C). To further quantify muscle damage, EBD, which penetrates damaged myofiber, was injected immediately after the Day 8 treadmill run (Fig. 3A), and the mice were sacrificed the following day. The TA muscle was then used to extract and quantify the concentration of EBD, assessing the extent of muscle damage. No statistically significant differences were found when analyzing all mice (Fig. 4D), as well as when analyzing mice that completed (Fig. 4E) or did not complete (Fig. 4F) the running separately. Taken together with the creatine phosphokinase data (Fig. 1I) and the results shown in Fig. 2D and E, which indicated significantly less damage in mice treated with 4.8 mg/ml GlcNAc without any forced exercise, these findings suggest that while GlcNAc does not prevent muscle damage formation induced by forced downhill exercises (repeated eccentric contractions), it exerts positive effects on the muscle health of *mdx* mice, particularly at a concentration of 4.8 mg/ml GlcNAc.

**Fig. 4.**
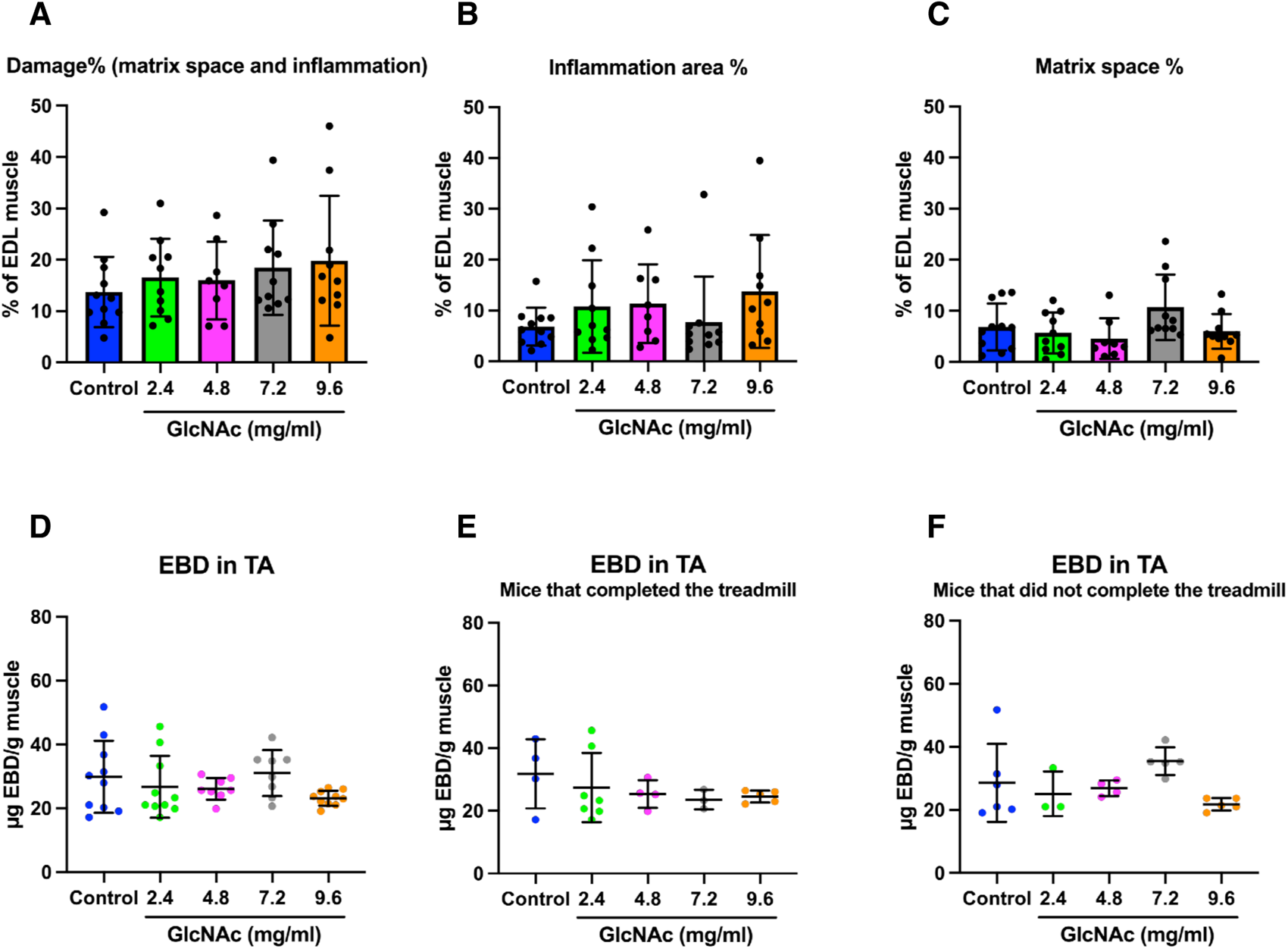
Effect of GlcNAc on Muscle Damage Formation in *mdx* Mice Subjected to Treadmill Running (Protocol 2) **A-C.** Tissue sections of the extensor digitorum longus (EDL) muscle were prepared and stained with hematoxylin and eosin (H&E). Images were captured and converted to grayscale for segmentation analysis using in-house software to distinguish between the inflammation area and matrix space. The percentage of the damaged area was calculated by combining the inflammation area and matrix space, normalized to the total area of the EDL (**A**). The percentage of the inflammation area (**B**) and matrix space (**C**) was calculated by normalizing these areas to the total EDL area. The sample size for panels **A-C** was 8 to 11. **D-F.** Muscles were stained with Evans Blue Dye (EBD), which specifically marks damaged muscle areas. The dye was then extracted to determine its concentration. Analysis was performed on all mice, including those that completed and did not complete the treadmill running on Day 8 (**D**), only those mice that completed the run (**E**), and only those that did not complete the run (**F**). The sample size for panel **D** was 10 to 11, and for panels **E** and **F**, it was 3 to 7. Statistical analysis was performed using one-way ANOVA with Tukey’s post-hoc test. No significant differences were observed. Data represent means ± standard deviations.

### Analysis of the Spontaneous Locomotor Activity of *mdx* Mice

The analysis of muscle damage through histological techniques can be affected by various factors, including the timing of damage formation. For example, significant damage may be detected if it occurs shortly before the sacrifice of a mouse, whereas earlier damage that has been partially repaired and regenerated may appear less severe. This variability is likely influenced by the autonomous behavior and continuous spontaneous locomotor activity of the mice. To minimize the influence of such ambiguous factors, it is essential to monitor the spontaneous locomotor activity of *mdx* mice continuously under conditions where no significant artificial stress is applied. For this purpose, we utilized the Digital Ventilated Cage (DVC^TM^) system, which employs 12 electrode sensors positioned at the bottom of the cage (Fig. 5A)(**39, 40**). Electromagnetic field lines are generated between the electrodes, and the presence of the mice disrupts these lines, allowing the system to detect their location and measure locomotor activity. This system allows for continuous, stress-free monitoring of spontaneous locomotor activity, providing insights into how GlcNAc affects the muscles involved in locomotion.

**Fig. 5.**
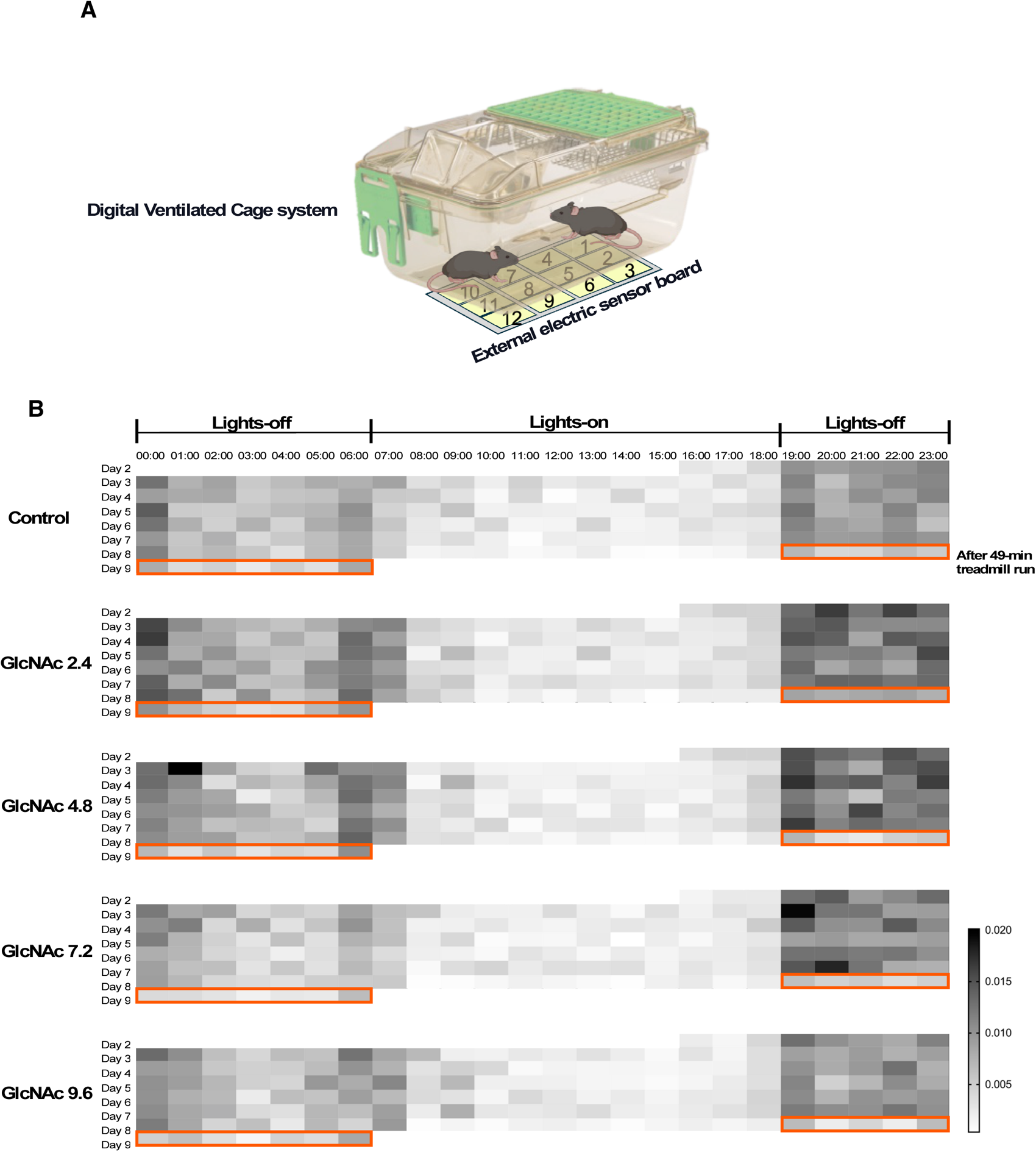
Overview of Locomotor Activity Analysis in *mdx* Mice (Protocol 2) **A.** The Digital Ventilated Cage (DVC) system was employed to monitor the locomotor activity of *mdx* mice. Twelve electrode sensors are positioned at the bottom of each cage, generating electromagnetic field lines between them. The presence of mice disrupts these field lines, enabling the system to detect their location and measure their locomotor activity. **B.** An example of locomotor activity measured over 6.5 days is illustrated. This example follows the schedule shown in Fig. 3A, which outlines the treadmill run. Locomotor activity measurement began on Day 2, the resting day after the first treadmill run. The activity is represented by a heatmap, with colors ranging from light gray (low activity) to dark gray (high activity). The data depicts hourly locomotor activity, with lights-off and lights-on periods marked. On Day 8, a 49-minute run was performed, and the orange box highlights the subsequent lights-off period, during which the mice’s locomotor activity was reduced.

The mouse was housed individually during the period of locomotor activity monitoring. Fig. 5B illustrates representative results of the average hourly locomotor activity of the mice, with higher activity levels shown in dark gray and lower activity levels in light gray on a heat map. During the lights-off period (12 hours), the mice exhibited higher locomotor activity, consistent with their nocturnal nature. The DVC system, therefore, accurately captures the natural circadian rhythm of the mice, reflecting their typical locomotor activity.

### Impact of GlcNAc on the Spontaneous Locomotor Activity of *mdx* Mice (Protocol 3)

In Protocol 3, we measured the spontaneous locomotor activity of *mdx* mice for one week following three weeks of GlcNAc treatment at concentrations of 2.4 to 9.6 mg/ml (Fig. 6A). Control data related to BW, survival, and the mass of TA, DEL, and Sol muscles are shown in Supplementary Fig. 1, with no significant differences observed. While Fig. 5B illustrates hourly average locomotor activity in a heat map format, Fig. 6A shows the cumulative locomotor activity over the monitoring period. The flat phase represents the locomotor activity during the lights-on period, with increased activity during the lights-off period (Fig. 6A). Among the *mdx* mice treated with GlcNAc, those receiving 2.4, 4.8, and 7.2 mg/ml showed significantly higher spontaneous locomotor activity compared to the control, suggesting that these doses of GlcNAc promote nocturnal locomotor activity in *mdx* mice. The activity of mice treated with 9.6 mg/ml was still higher than the control group, but lower than those treated with 2.4, 4.8, and 7.2 mg/ml. This finding suggests that there is an optimal GlcNAc concentration range for promoting locomotor activity.

**Fig. 6.**
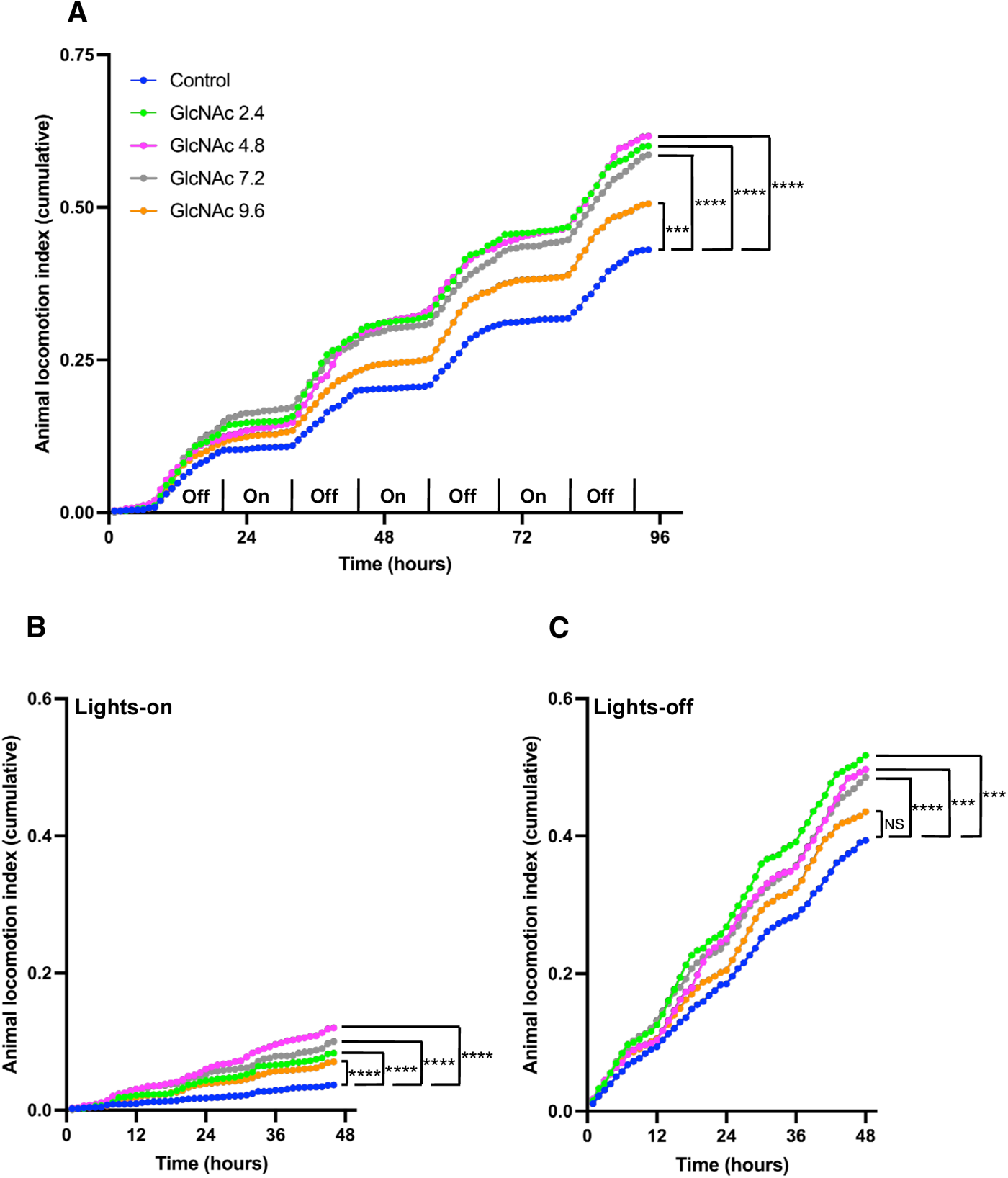
Effect of GlcNAc on the Locomotor Activity of *mdx* Mice (Protocol 3) Mice were treated with or without GlcNAc for 35 days, and locomotor activity was measured over the final 3.5 days before the end of the housing period. **A.** Locomotor activity during both lights-on and lights-off periods, as indicated in Frame **A**. **B.** Locomotor activity during the lights-on period. **C.** Locomotor activity during the lights-off period. Statistical analysis was performed using two-way ANOVA with Dunnett’s test. Significance levels are indicated as ***P < 0.001 and ****P < 0.0001, with “NS” denoting no significance. The number of mice used for Control, and for 2.4, 4.8, 7.2, and 9.6 mg/ml GlcNAc groups were 4, 5, 6, 5, and 6, respectively.

Fig. 6B and C present the locomotor activity during the lights-on and lights-off periods, respectively, with the results representing the cumulative activity during each corresponding period. During the lights-on period (Fig. 6B), *mdx* mice generally exhibited lower locomotor activity compared to the lights-off period (Fig. 6C). Nonetheless, GlcNAc treatment still promoted locomotor activity in *mdx* mice during both the lights-on and lights-off periods. Histological analysis of muscle damage (Fig. 2) suggests that only 4.8 mg/ml GlcNAc showed a significant effect in reducing damage formation. However, these locomotor activity results indicate that various doses of GlcNAc, particularly 2.4, 4.8, and 7.2 mg/ml, have a significant effect on enhancing the locomotor activity of *mdx* mice across different times of the day. Additionally, these doses of GlcNAc did not show any effect on BW, survival rate, or the mass of TA, EDL, and Sol muscles (supplemental Fig. 1), suggesting that the promoted locomotor activity did not have any harmful effects on the *mdx* mice.

### Impact of GlcNAc on the Spontaneous Locomotor Activity of *mdx* Mice Subjected to Treadmill Running (Protocol 2)

In Protocol 2, we analyzed the locomotor activity of *md*x mice subjected to treadmill running to assess the effects of GlcNAc under exercise conditions. The monitoring period lasted about 6 days, during which the mice underwent the second and third acclimation treadmill runs at speeds of 10 and 15 m/min for 10 minutes on Day 3 and Day 5, respectively, during the lights-on period, as illustrated in Fig. 3A. Following the acclimation phase, a downhill treadmill run with a gradual increase in speed from 8 to 15 m/min over a total of 49 minutes at a 15-degree incline was performed. DVC analysis showed that the effects of 7.2 and 9.6 mg/ml GlcNAc in mice subjected to forced exercise reduced spontaneous locomotor activity to levels similar to the control group (Fig. 6A vs Fig. 7A), suggesting that forced exercise diminishes the positive impact of these doses on activity. In contrast, *mdx* mice treated with 2.4 and 4.8 mg/ml GlcNAc still exhibited significantly higher locomotor activity compared to the control group as observed in mice without forced exercise. During the lights-on period, only the 2.4 mg/ml GlcNAc-treated group showed significantly higher locomotor activity, while the other groups displayed similar or slightly reduced activity relative to the control (Fig. 7B). Notably, during the lights-off period, *mdx* mice treated with 2.4 and 4.8 mg/ml GlcNAc showed significantly higher locomotor activity (Fig. 7C), suggesting that these doses of GlcNAc positively impact muscle health, enhancing resistance to consecutive exercises, and supporting muscle movement even after repeated eccentric contractions.

**Fig. 7.**
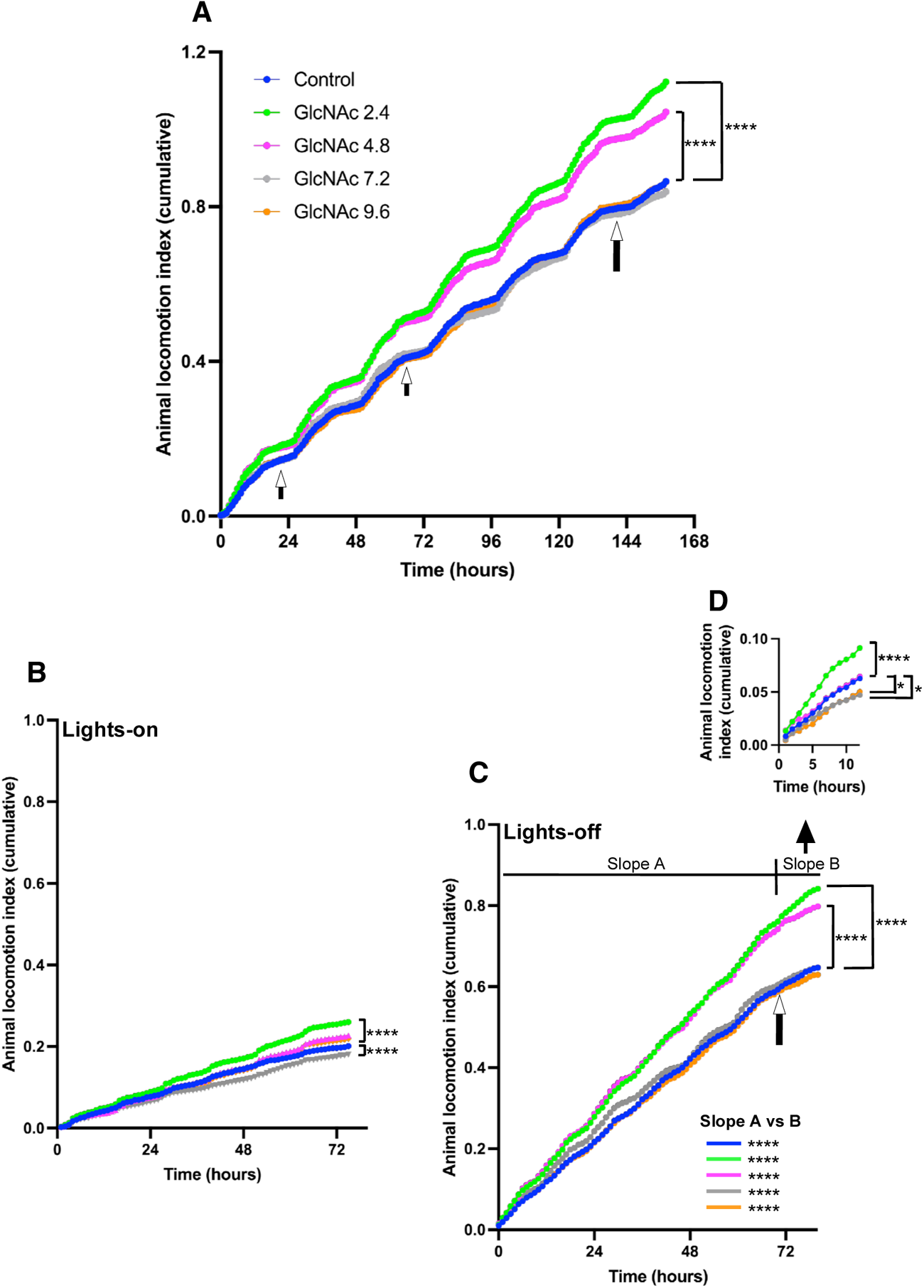
Effect of GlcNAc on the Locomotor Activity of *mdx* Mice Subjected to Treadmill Running (Protocol 2) Mice were treated with or without GlcNAc for 35 days. In the last week of the treatment, mice were subject to treadmill running according to the schedule illustrated in Fig. 3A. Locomotor activity was measured over the final 6.5 days before the mice were sacrificed. **A.** Locomotor activity during both lights-on and lights-off periods. The first and second small arrows correspond to the acclimation treadmill runs performed on Day 3 and Day 5, respectively (Fig. 3A). The large arrow indicates when the 49-minute treadmill run with a 15-degree downhill incline, gradually increasing in speed from 8 to 15 m/min, was conducted. **B.** Locomotor activity during the lights-on period. **C.** Locomotor activity during the lights-off period. The arrow marks the time of the 49-minute treadmill run. “Slope A” and “Slope B” refer to the periods before and after the run, respectively. **D.** Cumulative locomotor activity after the 49-minute treadmill run. Statistical analysis was performed using two-way ANOVA with Dunnett’s test. For the analysis of Slope A and Slope B, two-way ANOVA with Tukey’s test was used. Significance levels are indicated as *P < 0.05 and ****P < 0.0001. The number of mice used in the Control group and the 2.4, 4.8, 7.2, and 9.6 mg/ml GlcNAc groups were 12, 12, 10, 10, and 10, respectively.

The heat map shown in Fig. 5B corresponds to the *mdx* mice subjected to treadmill running (Protocol 2). After the 49-minute treadmill run performed on Day 8 during the lights-on period, the locomotor activity of all groups of *mdx* mice remained relatively low, even during the lights-off period (Fig. 5B, Day 8). This consistency likely reflects the muscle damage induced by the 49-minute treadmill run, as demonstrated in Fig. 4. In Fig. 7C, the locomotor activity before (Slope A) and after (Slope B) the 49-minute run was analyzed, and in all groups, the activity was statistically significantly reduced, suggesting the negative impact of repeated excentric contractions. Cumulative locomotor activity after the 45-minute run is shown separately in Fig. 7D. Despite of muscle damages induced by downhill running, mice treated with 2.4 mg/ml retained significantly higher locomotor activity compared to the control group.

These results, combined with the locomotor activity data, shows that 2.4 and 4.8 mg/ml GlcNAc can promote spontaneous locomotor activity in mice subjected to the acclimation phase level of exercise (horizontal treadmill running) (Slope A). However, there appears to be a threshold of exercise intensity beyond which 4.8 mg/ml GlcNAc becomes ineffective. Additionally, it is important to note that even though GlcNAc becomes ineffective beyond this threshold, it does not appear to have any harmful effects on *mdx* mice, as no significant impact on the mice’s BW, survival, or muscle mass was found (Fig. 3).

### Effect of GlcNAc and Prednisolone on the Locomotor Activity of *mdx* Mice (Protocol 3)

Prednisolone is commonly used in the intervention of DMD patients to suppress inflammation (10–12). In Protocol 3, we also investigated how prednisolone influences the effects of GlcNAc. As shown in Fig. 8, the combination treatment of prednisolone with various doses of GlcNAc did not significantly affect BW (Fig. 8A), survival (Fig. 8B), TA muscle mass (Fig. 8C and D), EDL muscle mass (Fig. 8E and F), or Sol muscle mass (Fig. 8G and H). We then used the DVC system to evaluate the locomotor activity under combination treatment with prednisolone and GlcNAc. As expected, prednisolone alone significantly increased the spontaneous locomotor activity of *mdx* mice compared to the control group (Fig. 9A). Among the GlcNAc-treated groups, only the 2.4 mg/ml GlcNAc combined with prednisolone showed a statistically significant increase in overall locomotor activity compared to prednisone alone.

**Fig. 8.**
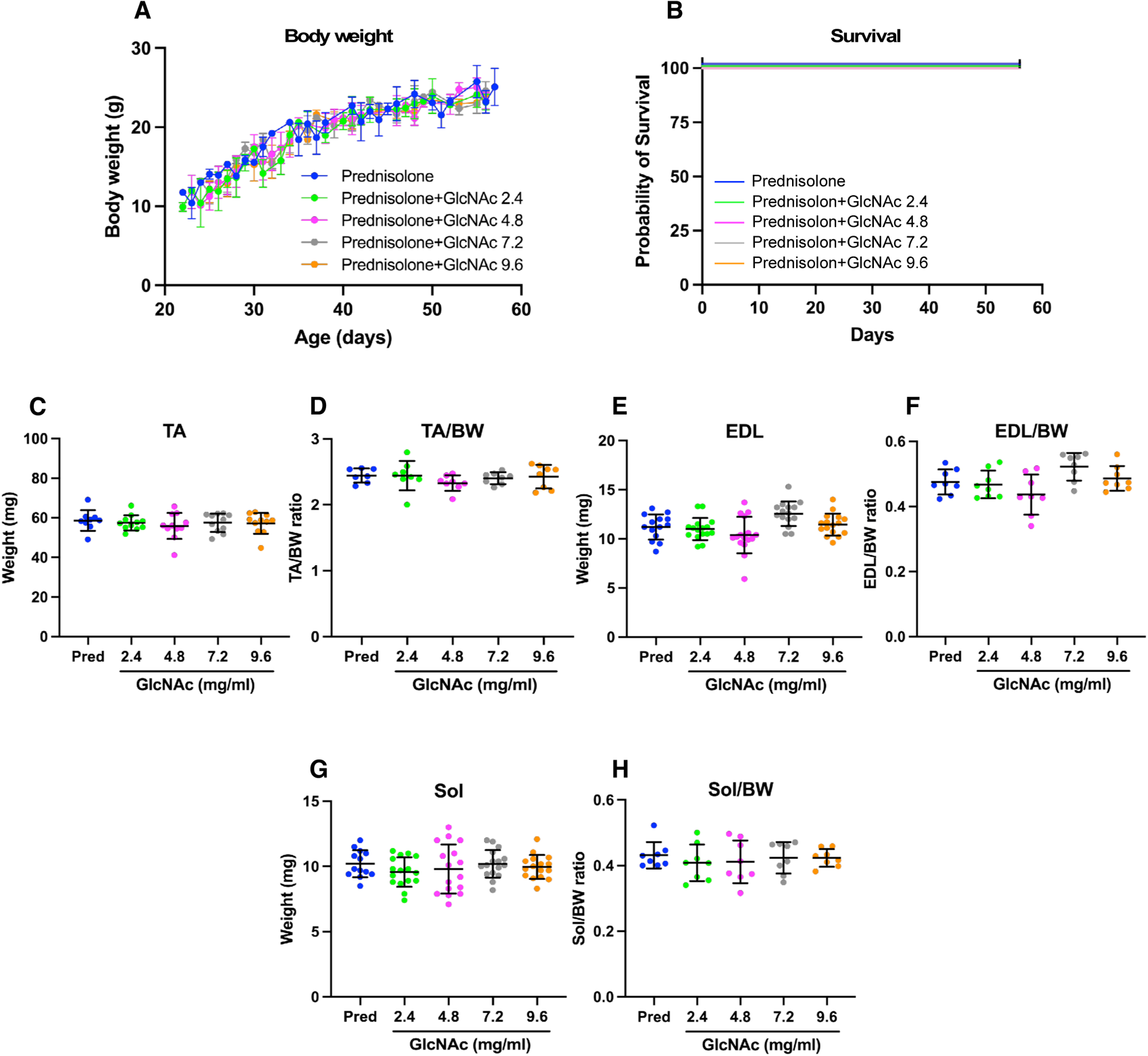
Effect of GlcNAc on Muscle Damage Formation in *mdx* Mice treated with Prednisolone (Protocol 3) GlcNAc (2.4, 4.8, 7.2, and 9.6 mg/ml) was administered orally along with or without 1 mg/kg body weight (BW) per day of prednisolone to *mdx* mice via voluntary intake. **A.** Mice were treated for 35 days, during which their BW was regularly measured. **B.** The survival rate of the mice was monitored throughout the study. For both **A** and **B**, the number of mice used was 8. **C-H.** After 35 days of treatment, the *mdx* mice were sacrificed, and the mass of the tibialis anterior (TA) (**C**), TA mass relative to BW (**D**), extensor digitorum longus (EDL) mass (**E**), EDL mass relative to BW (**F**), soleus (Sol) mass (**G**), and Sol mass relative to BW (**H**) were measured. For **C, E** and **G**, the sample size was 16, and for **D, F** and **H,** the sample size was 8. Statistical analyses were performed using ordinary one-way ANOVA with Tukey’s post-hoc test (**A**, and **C-I)**, and Mantel-Cox test (**B**). No significant differences were observed. **A**, and **C-I**. Data represent means ± standard deviations.

**Fig. 9.**
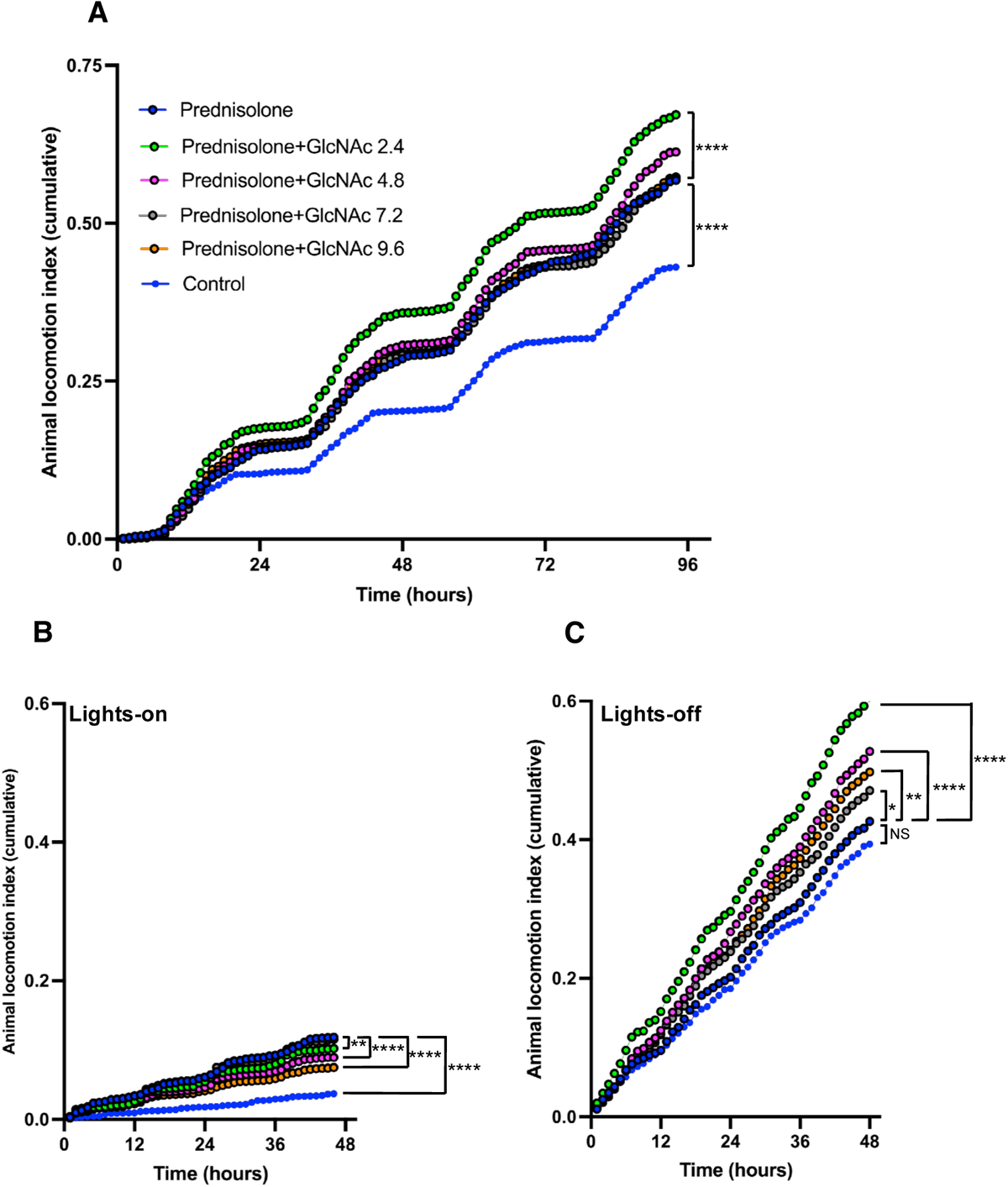
Effect of GlcNAc and Prednisolone on the Locomotor Activity of *mdx* Mice (Protocol 3) Mice were treated for 35 days and the locomotor activity was measured over the final 3.5 days before the end of the treatment period. **A.** Locomotor activity during both lights-on and lights-off periods. **B.** Locomotor activity during the lights-on period. **C.** Locomotor activity during the lights-off period. Statistical analysis was performed using two-way ANOVA with Dunnett’s test. Significance levels are indicated as *P < 0.05, ***P < 0.001, and ****P < 0.0001, with “NS” denoting no significance. The number of mice used for Control, and for 2.4, 4.8, 7.2, and 9.6 mg/ml GlcNAc groups were 6, 8, 8, 8, and 8, respectively.

Further analysis of locomotor activity during the lights-on period (Fig. 9B) revealed that prednisolone alone significantly increased locomotor activity, whereas GlcNAc mitigated this increase. In contrast, during the lights-off period, all doses of GlcNAc combined with prednisolone enhanced the locomotor activity of *mdx* mice compared to treatment with prednisolone alone (Fig. 9C), suggesting GlcNAc has the capacity to further improve muscle health of mice undergoing prednisolone treatment. Due to GlcNAc’s counteracting effect on prednisolone during the lights-on period and its promotion of activity during the lights-off period, only the 2.4 mg/ml GlcNAc dose showed a significant effect when considering combined lights-on and lights-off data (Fig. 9A). Prednisolone is known to have many side effects including insomnia (41), although there were few reports examining its effects on insomnia in mdx mice. As shown in Fig. 9B, the significant increase in abnormal nocturnal activity in mice treated with prednisolone appears to be associated with insomnia. These results suggest that GlcNAc may counteract this abnormal activity during the lights-on period. Nevertheless, these results suggest that the combination of GlcNAc and prednisolone promotes nocturnal locomotor activity during the lights-off period. Given that the combination treatment did not impact BW, survival, or muscle mass (Fig. 8), it is unlikely to produce adverse effects. Table 1 summarizes the results, showing that GlcNAc, particularly at 2.4 mg/ml, consistently promotes the locomotor activity of *mdx* mice.

**Table 1.** Summary of the results of locomotor activity. Fold difference relative to Control (water) and prednisolone are calculated. Results obtained using 2.4 mg/ml GlcNAc were indicated by black and right blue bold characters.

| Treatment | Lights | locomotor activity | Relative to Control (fold) | Relative to Prednisolone (fold) |
| --- | --- | --- | --- | --- |
| Control | On and Off | 0.20240 |  |  |
| **GlcNAc 2.4** | **On and Off** | **0.29610** | **1.46294** |  |
| GlcNAc 4.8 | On and Off | 0.29400 | 1.45257 |  |
| GlcNAc 7.2 | On and Off | 0.28750 | 1.42045 |  |
| GlcNAc 9.6 | On and Off | 0.24430 | 1.20702 |  |
| Control | On | 0.01860 |  |  |
| **GlcNAc 2.4** | **On** | **0.04060** | **2.18280** |  |
| GlcNAc 4.8 | On | 0.05900 | 3.17204 |  |
| GlcNAc 7.2 | On | 0.05000 | 2.68817 |  |
| GlcNAc 9.6 | On | 0.03150 | 1.69355 |  |
| Control | Off | 0.20070 |  |  |
| **GlcNAc 2.4** | **Off** | **0.27930** | **1.39163** |  |
| GlcNAc 4.8 | Off | 0.25580 | 1.27454 |  |
| GlcNAc 7.2 | Off | 0.25800 | 1.28550 |  |
| GlcNAc 9.6 | Off | 0.23020 | 1.14699 |  |
| Control+Treadmill | On and Off | 0.46640 |  |  |
| **GlcNAc 2.4+Treadmill** | **On and Off** | **0.59020** | **1.26544** |  |
| GlcNAc 4.8+Treadmill | On and Off | 0.56530 | 1.21205 |  |
| GlcNAc 7.8+Treadmill | On and Off | 0.46270 | 0.99207 |  |
| GlcNAc 9.6+Treadmill | On and Off | 0.46130 | 0.98907 |  |
| Control+Treadmill | On | 0.11460 |  |  |
| **GlcNAc 2.4+Treadmill** | **On** | **0.14080** | **1.22862** |  |
| GlcNAc 4.8+Treadmill | On | 0.12100 | 1.05585 |  |
| GlcNAc 7.8+Treadmill | On | 0.09770 | 0.85253 |  |
| GlcNAc 9.6+Treadmill | On | 0.11620 | 1.01396 |  |
| Control+Treadmill | Off | 0.36050 |  |  |
| **GlcNAc 2.4+Treadmill** | **Off** | **0.46160** | **1.28044** |  |
| GlcNAc 4.8+Treadmill | Off | 0.45500 | 1.26214 |  |
| GlcNAc 7.8+Treadmill | Off | 0.37510 | 1.04050 |  |
| GlcNAc 9.6+Treadmill | Off | 0.35400 | 0.98197 |  |
| Control | On and Off | 0.20240 |  | 0.73360 |
| Prednisolone | On and Off | 0.27590 | 1.36314 |  |
| **GlcNAc 2.4** | **On and Off** | **0.33500** | **1.65514** | **1.21421** |
| GlcNAc 4.8 | On and Off | 0.29030 | 1.43429 | 1.05219 |
| GlcNAc 7.8 | On and Off | 0.27340 | 1.35079 | 0.99094 |
| GlcNAc 9.6 | On and Off | 0.27820 | 1.37451 | 1.00834 |
| Control | On | 0.01960 |  | 0.29878 |
| Prednisolone | On | 0.06560 | 3.34694 |  |
| **GlcNAc 2.4** | **On** | **0.05680** | **2.89796** | **0.86585** |
| GlcNAc 4.8 | On | 0.05020 | 2.56122 | 0.76524 |
| GlcNAc 7.8 | On | 0.06450 | 3.29082 | 0.98323 |
| GlcNAc 9.6 | On | 0.04310 | 2.19898 | 0.65701 |
| Control | Off | 0.20070 |  | 0.91854 |
| Prednisolone | Off | 0.21850 | 1.08869 |  |
| **GlcNAc 2.4** | **Off** | **0.31540** | **1.57150** | **1.44348** |
| GlcNAc 4.8 | Off | 0.27030 | 1.34679 | 1.23710 |
| GlcNAc 7.8 | Off | 0.24560 | 1.22372 | 1.12403 |
| GlcNAc 9.6 | Off | 0.25570 | 1.27404 | 1.17025 |

## Discussion

GlcNAc, an endogenous precursor of UDP-GlcNAc, serves as a substrate for various N-acetylglucosaminyltransferases. Importantly, its intracellular concentration is closely associated with the levels of acetyllactosamine-rich N-linked oligosaccharides, which are recognized by the oligosaccharide-binding protein Gal-3 (13–25). This association is particularly significant because the rate-limiting enzyme for the biosynthesis of these oligosaccharides requires a high concentration of UDP-GlcNAc (30, 42, 43). Notably, the interaction between Gal-3 and acetyllactosamine-rich N-linked oligosaccharides plays a critical role in regulating the lateral movement of various membrane proteins (dynamics), as well as mediating cell-to-cell and cell-to-extracellular matrix interactions (17–22). On the cell surface, these oligosaccharides contribute not only to the formation of the glycocalyx, which extends 50-500 nm above the cell surface but also to determining cell characteristics by presenting specific concentrations and aggregations of acetyllactosamine residues (17–22). The glycocalyx is vital for facilitating flexible interactions that support dynamic cellular processes (44). Altering the glycocalyx by increasing the synthesis of acetyllactosamine-rich N-linked oligosaccharides through GlcNAc supplementation can potentially influence cell dynamics governed by the glycocalyx (17–22).

Our previous and recent research demonstrated that GlcNAc promotes myotube formation in C2C12 cells and primary myoblasts(13, 25), and intraperitoneal administration of GlcNAc in *mdx* mice enhances muscle force (13)These findings suggest that GlcNAc changes in the nature of the glycocalyx in the muscles of *mdx* mice, although the full implications of these changes remained unclear. In this study, we demonstrated that orally administered GlcNAc improves overall muscle health and significantly promotes the spontaneous locomotor activity of *mdx* mice.

We tested the effects of different GlcNAc doses dissolved in the drinking water at the concentrations of 2.4, 4.8, 7.2, and 9.6 mg/ml, which was equivalent to 0.6, 1.2, 1.8, and 2.4 g/kg BW per day, respectively, based on the observed mice’s daily water intake. These doses in mice are equivalent to doses of 72, 144, 216, and 288 mg/kg BW per day, respectively, in humans, based on the body surface area normalization method (34). Previous clinical studies for patients with inflammatory bowel disease used 6 and 12 g GlcNAc per patient with the analysis revealing the therapeutical benefit (36). The doses used in this study is relatively close to or slightly higher than the one in those studies and several doses of GlcNAc consistently exhibit therapeutical impact, including spontaneous locomotor activity, and reduced muscle damage. In our previous study, 250 mg/kg BW of GlcNAc per day for intraperitoneal administration for 10 days exhibit the mitigation of DMD progression, including reduced muscle damage(13). Considering the established efficient absorption of orally administered GlcNAc (30), it is possible that doses lower than 2.4 mg/ml (600 mg/kg BW) GlcNAc may be sufficient for therapeutic effects. Therefore, it is essential to determine the minimal and most optimal doses for the preparation of a future clinical study in individuals with DMD.

Our findings indicated that GlcNAc promotes spontaneous locomotor activity in *mdx* mice. This promotion likely indicates an improvement in the muscular health of *mdx* mice, though alternative explanations, such as potential adverse effects, cannot be entirely ruled out. However, our examination revealed no apparent effects of GlcNAc on BW, survival, or the mass of TA, EDL, and Sol muscles. Previous preclinical chronic toxicity studies demonstrated that oral administration of GlcNAc 2.5 g/kg BW per day (equivalent to 0.6 g/kg BW per day in humans) for 52 weeks results in no adverse effects or histopathological changes on tissues (32, 33), suggesting that the promotive effect on spontaneous locomotor activity is not caused by any adverse effects, and likely improves the muscular health of the mice. While most of GlcNAc doses in this study appear to promote locomotor activity, GlcNAc at 2.4 and 4.8 mg/ml showed higher promoting effects than 7.8 and 9.6 mg/ml of GlcNAc, suggesting that an optimal dose likely exists.

Our previous reports indicated that intraperitoneal GlcNAc administration improves *ex vivo* muscular force in *mdx* mice (13), suggesting that GlcNAc alters the quality, rather than the quantity, of muscles. The promoted locomotor activity observed in orally GlcNAc-treated mice is likely related to these qualitative changes in muscles. This change may help prevent muscles from sustaining damage. To investigate this, we analyzed sections of EDL muscles and quantified the inflammatory area and matrix space, finding a significant reduction in damage formation with 4.8 mg/ml GlcNAc administration. However, the extent of damage varied significantly, with some damage appearing to have formed immediately before sacrifice and others over a time frame that allowed some regeneration. This variability made it challenging to assess whether other GlcNAc doses reduced muscular damage. To eliminate such experimental ambiguity and directly address whether GlcNAc protects skeletal muscles from repeated eccentric contractions, we conducted a downhill treadmill run to induce muscular damage one day before sacrifice. Our results indicated that GlcNAc does not prevent damage formation induced by repeated eccentric contractions, suggesting that the observed reduction in damage with 4.8 mg/ml GlcNAc may result from promoting muscle fiber regeneration, consistent with our *in vitro* observations that GlcNAc promotes myogenesis (13, 25). In our previous report, *ex vivo* contractile property analysis of EDL and Sol muscle of mice administrated GlcNAc intraperitoneal shows an increase in the maximum specific force production of the EDL and Sol, while no protection was observed from *ex vivo* repeated eccentric contraction stimuli (13), which is also consistent with these results.

Regarding the spontaneous locomotor activity of *mdx* mice subjected to treadmill running, 2.4 and 4.8 mg/ml GlcNAc promoted locomotor activity during the acclimated running phase (horizontal running), whereas 7.8 and 9.6 mg/ml did not show any promoting effects. The lack of positive impact from the 7.8 and 9.6 mg/ml doses, which showed therapeutic effects when mice were not subject to forced exercises, suggests that a series of horizontal treadmill exercises, even without repeated eccentric contractions is sufficient to negatively impact skeletal muscle. In contrast, 2.4 and 4.8 mg/ml doses showed the promoted locomotor activity. Considering that muscle regeneration and the formation of myofibers require at least 10 days (45), the lack of impact on locomotor activity during the period of focused exercises is likely independent of new myogenesis of the damaged myofibers and is instead related to the overall muscle health and resistance to exercise-induced damage. In other words, this resistance is closely linked to GlcNAc-promoted muscle health during the first 3 weeks of administration. Indeed, although all GlcNAc treatments increased the spontaneous locomotor activity of mice that were not subject to forced exercises, the activity level with the 9.6 mg/ml treatment was significantly lower than that with 2.4 mg/ml (p<0.1). This implies that, even though this highest dose exhibited a therapeutic potential to increase locomotor activity, it failed to improve muscle health to the extent that it could resist damage induced by horizontal treadmill exercise. When mice were subjected to 49 minutes of downhill treadmill running, involving repeated eccentric contractions, only 2.4 mg/ml treatment exhibited increased locomotor activity, while other doses did not. This suggests that appropriate control of muscular damage formation may be essential for making GlcNAc effective on muscle functions.

Regenerating muscle fibers requires a proper extracellular matrix structure to guide fiber regeneration (45, 46). Events that disturb this structure, including excessive inflammation, could negatively impact regeneration efficiency. Corticosteroids, such as prednisolone, which have anti-inflammatory effects, are therefore used to slow the progression of muscular dystrophy (10–12). We studied the effect of GlcNAc on *mdx* mice treated with prednisolone. As expected, prednisolone alone promoted spontaneous locomotor activity in *mdx* mice, indicating an improvement in muscle health. GlcNAc further enhanced the locomotor activity, though the effects were complex. During the lights-off period, GlcNAc significantly increased nocturnal spontaneous locomotor activity relative to mice treated with prednisolone alone, suggesting that GlcNAc has the potential to improve the clinical status of prednisolone-treated DMD patients. Conversely, during the lights-on period, GlcNAc reduced the locomotor activity, which was increased by prednisolone compared to the control. One of the side effects of prednisolone is insomnia (41), which may explain why abnormal nocturnal spontaneous locomotion during the lights-on period was significantly higher in prednisolone-treated mice compared to the control, a difference that the DVC system was sensitive enough to detect. GlcNAc treatment significantly reduced this prednisolone-induced hyperactivity. The exact reason remains unknown but it appears that GlcNAc counteracts this hyperreactive response in mice treated with prednisolone.

We recently demonstrated that GlcNAc can create an optimal environment for myogenesis by regulating myoblasts’ coordinated flow and aligning myoblasts along the eventual shapes of regenerated myofibers (25). Together with this study, GlcNAc likely exerts its effects by promoting myofiber regeneration rather than preventing damage formation, leading to the promotion of locomotor activity in *mdx* mice. Given that GlcNAc does not show any adverse effects, it may be used to improve the muscle status of DMD patients. Additionally, the GlcNAc concentration in the blood has been shown to decrease with aging in wild-type and *mdx* mice (Supplementary Fig. 2) (47), suggesting GlcNAc supplementing could be beneficial in this context. While the optimal doses of GlcNAc for improving human muscle status still need to be determined, this current study suggests that GlcNAc has the potential to improve muscle status alone or in combination with other drugs, such as corticosteroids.

## Materials and Methods

### Animal Studies

All animal experiments were conducted in accordance with the policies of the Comités de Protection des Animaux at Université Laval (CPAUL-3). Male *mdx* dystrophic mice (C57BL/10ScSn-Dmd^mdx^/J) were purchased from The Jackson Laboratory (Bar Harbor, ME, USA) and bred at the CRCHU animal facility. The mice were housed under controlled conditions: temperature (23°C ± 2°C), humidity (50% ± 5%), and light (12:12 hour light-dark cycle). They had unlimited access to food and water. GlcNAc (2.4, 4.8, 7.2, and 9.6 mg/ml) and/or prednisolone (1 mg/kg BW per day, based on a 5 ml daily water intake) were administered orally via voluntary intake to the *mdx* mice. All treatments started at the age of 3 weeks and the treatment lasted for 35 ± 1 days.

### Protocols

**Protocol 1:** GlcNAc was orally administered through drinking water at doses of 0, 2.4, 4.8, and 9.6 mg/ml for 35 days. Mice were sacrificed for the examination of muscles and serum CPK levels.

**Protocol 2:** GlcNAc was orally administered through drinking water at doses of 0, 2.4, 4.8, 7.2, and 9.6 mg/ml for the entire 35 days. After 26 ± 1 days of treatment, the mice were subject to treadmill exercises (see below for the detailed conditions, Fig. 3A). Spontaneous locomotor activity for the last 6.5 days was monitored using the DVC system. Twenty-four hours before their sacrifice, mice were injected with EBD.

**Protocol 3:** Mice were orally administered GlcNAc at doses of 0, 2.4, 4.8, 7.2, and 9.6 mg/ml, with or without prednisolone for 35 ± 1 days. Spontaneous locomotor activity for the last 3.5 days was monitored using the DVC system.

### Materials

GlcNAc (100% pure) was obtained from Wellesley Therapeutics, Ontario, Canada. Prednisolone was purchased from Abcam.

### CPK Activity Assay

Blood samples were centrifuged to separate the supernatant after incubation at room temperature. Serum CPK assays were performed using the Pointe Scientific Creatine Kinase Liquid Reagent Set, following the manufacturer’s protocol. Serum (5 μl) was incubated with 100 μl of the working reagent at 37°C. Dye absorbance was measured spectrophotometrically at 340 nm over a period of 5 minutes. The values were calculated according to the manufacturer’s instructions, and the data were expressed as units per liter (CPK U/L).

### Muscle EVD Uptake Experiments

Muscle damage was assessed using the penetration of EBD (Sigma-Aldrich) into muscles following the method described by Wooddell et al. (47). Briefly, EBD was dissolved in phosphate-buffered saline at a concentration of 10 mg/ml and injected intraperitoneally at a dose of 100 μl per 10 g of BW. Twenty-four hours after the injection, the mice were sacrificed, and the TA muscles were harvested and frozen at -80°C. The frozen TA muscles were then ground into a powder, and 50 mg of the powder was used for quantification. *N*,*N*-Dimethylformamide (1 ml) was added to the muscle powder, followed by incubation for 24 hours at room temperature. After centrifugation at 6,500 g for 25 minutes, the supernatant was collected, and the absorbance was measured at 630 nm to evaluate the extent of muscle damage.

### Histological Analysis

Muscles were dissected and frozen in Tissue-Plus™ O.C.T. Compound (Fisher Healthcare) using isopentane cooled with liquid nitrogen. Frozen muscle sections (8–10 μm) were fixed in cold acetone or 4% paraformaldehyde. H&E staining was performed following standard protocols. Images of the H&E-stained muscle sections were captured using a standard microscope equipped with a color camera. The images were then converted to grayscale and segmented using in-house software available on GitHub (DOI 10.5281/zenodo.12988726) (48). Segmented images were used to quantify muscle area, muscular fiber size, inflammatory area, and matrix space. Quantification was performed with manual correction as needed. The percentages of inflammatory area and matrix space were then calculated.

### Treadmill Performance Tests

The treadmill performance test was conducted using a motor-driven MK-680S treadmill system (Columbus Instruments) to evaluate the muscle performance of mice after 3 weeks of housing. On Day 1, the mice were placed on the treadmill and ran for 10 minutes at a speed of 5 m/min on a flat surface. After a rest day on Day 2, the mice ran again on Day 3 for 10 minutes at an increased speed of 10 m/min. Following another rest day on Day 4, a third run was performed on Day 5 for 10 minutes at a speed of 15 m/min on a flat treadmill. After two more rest days on Day 6 and Day 7, the final treadmill run was conducted on Day 8 on a 15-degree downhill incline, following this protocol: 2 minutes at 8 m/min, 2 minutes at 10 m/min, 5 minutes at 12 m/min, 5 minutes at 13 m/min, and 5 minutes at 14 m/min. During the last 30 minutes, the run was conducted at 15 m/min, for a total duration of 49 minutes. If a mouse stopped running, it was given a 2-minute rest before resuming the downhill treadmill run. If the same mouse stopped twice, it was returned to its cage.

### Locomotor Activity Measurement

The locomotor activity of mice was measured using the DVC system (Tecniplast, Italy), which is equipped with 12 electrodes. One mouse was housed in each cage under a 12:12-hour light: dark cycle. Locomotor activity was continuously monitored, and the data were stored using the DVC system’s associated data storage. The collected data were then analyzed using Prism GraphPad Software.

### Statistical analysis

Statistical significance was determined with one-way ANOVA and two-way ANOVA (the significances are explained in the figure legends). All statistical analyses were performed with Prism (GraphPad Software, La Jolla, CA, USA), and differences were considered significant at P < 0.05.

## Data availability

All data are included in the article and/or supporting information. The software was deposited to GitHub; DOI 10.5281/zenodo.12988726. Any questions regarding data availability can be directed to the corresponding author.

## Acknowledgments

We thank Willem Wassenaar (Wellesley Therapeutics, Ontario) for providing GlcNAc and the Bioimaging Platform of the Research Centre for Infectious Diseases, Axe of Infectious and Inflammatory Diseases at the CHU de Québec Research Centre, for their invaluable technical support with the microscopes. We also extend our gratitude to Giorgio Rosati, Senior Product Manager at DIGILAB, Tecniplast, for his technical advice and support. This research was financially supported by the Canada Foundation for Innovation and the Canadian Institutes of Health Research.

## Author contributions

M.S.S. and S.S. designed the study, conducted experiments, and contributed to manuscript’s writing. G.S-P and M.F. conducted experiments. A.R. conducted experiments and analyzed H&E staining data.

## References

1. Duan, D., Goemans, N., Takeda, S., Mercuri, E., and Aartsma-Rus, A. (2021) Duchenne muscular dystrophy. Nat. Rev. Dis. Prim. 7, 13

2. Roberts, T. C., Wood, M. J. A., and Davies, K. E. (2023) Therapeutic approaches for Duchenne muscular dystrophy. Nat. Rev. Drug Discov. 22, 917–934

3. Hoffman, E. P., Brown, R. H., and Kunkel, L. M. (1987) Dystrophin: The protein product of the duchenne muscular dystrophy locus. Cell. 51, 919–928

4. Fairclough, R. J., Wood, M. J., and Davies, K. E. (2013) Therapy for Duchenne muscular dystrophy: renewed optimism from genetic approaches. Nature reviews. Genetics. 14, 373–378

5. Ibraghimov-Beskrovnaya, O., Ervasti, J. M., Leveille, C. J., Slaughter, C. A., Sernett, S. W., and Campbell, K. P. (1992) Primary structure of dystrophin-associated glycoproteins linking dystrophin to the extracellular matrix. Nature. 355, 696–702

6. Koenig, M., Monaco, A. P., and Kunkel, L. M. (1988) The complete sequence of dystrophin predicts a rod-shaped cytoskeletal protein. Cell. 53, 219–228

7. Ervasti, J. M., Ohlendieck, K., Kahl, S. D., Gaver, M. G., and Campbell, K. P. (1990) Deficiency of a glycoprotein component of the dystrophin complex in dystrophic muscle. Nature. 345, 315–319

8. Ohlendieck, K., and Campbell, K. P. (1991) Dystrophin-associated proteins are greatly reduced in skeletal muscle from mdx mice. J. cell Biol. 115, 1685–1694

9. Rader, E. P., Turk, R., Willer, T., Beltrán, D., Inamori, K., Peterson, T. A., Engle, J., Prouty, S., Matsumura, K., Saito, F., Anderson, M. E., and Campbell, K. P. (2016) Role of dystroglycan in limiting contraction-induced injury to the sarcomeric cytoskeleton of mature skeletal muscle. Proc. Natl. Acad. Sci. 113, 10992–10997

10. Gloss, D., Moxley, R. T., Ashwal, S., and Oskoui, M. (2016) Practice guideline update summary: Corticosteroid treatment of Duchenne muscular dystrophy Report of the Guideline Development Subcommittee of the American Academy of Neurology. Neurology. 86, 465–472

11. Czifrus, E., and Berlau, D. J. (2024) Corticosteroids for the treatment of duchenne muscular dystrophy: a safety review. Expert Opin. Drug Saf. 10.1080/14740338.2024.2394578

12. Guglieri, M., Clemens, P. R., Perlman, S. J., Smith, E. C., Horrocks, I., Finkel, R. S., Mah, J. K., Deconinck, N., Goemans, N., Haberlova, J., Straub, V., Mengle-Gaw, L. J., Schwartz, B. D., Harper, A. D., Shieh, P. B., Waele, L. D., Castro, D., Yang, M. L., Ryan, M. M., McDonald, C. M., Tulinius, M., Webster, R., McMillan, H. J., Kuntz, N. L., Rao, V. K., Baranello, G., Spinty, S., Childs, A.-M., Sbrocchi, A. M., Selby, K. A., Monduy, M., Nevo, Y., Vilchez-Padilla, J. J., Nascimento-Osorio, A., Niks, E. H., Groot, I. J. M. de Katsalouli, M., James, M. K., Anker, J. van den Damsker, J. M., Ahmet, A., Ward, L. M., Jaros, M., Shale, P., Dang, U. J., and Hoffman, E. P. (2022) Efficacy and Safety of Vamorolone vs Placebo and Prednisone Among Boys With Duchenne Muscular Dystrophy. JAMA Neurol. 79, 1005–1014

13. Rancourt, A., Dufresne, S. S., St-Pierre, G., Levesque, J.-C., Nakamura, H., Kikuchi, Y., Satoh, M. S., Frenette, J., and Sato, S. (2018) Galectin-3 and N-acetylglucosamine promote myogenesis and improve skeletal muscle function in the mdx model of Duchenne muscular dystrophy. FASEB journal : official publication of the Federation of American Societies for Experimental Biology. 10.1096/fj.201701151rrr

14. Stanley, P., Moremen, K. W., Lewis, N. E., Taniguchi, N., and Aebi, M. (2022) N-Glycans. 10.1101/glycobiology.4e.9

15. Kizuka, Y. (2024) Regulation of intracellular activity of N-glycan branching enzymes in mammals. J. Biol. Chem. 300, 107471

16. Cummings, R. D., Liu, F.-T., Rabinovich, G. A., Stowell, S. R., and Vasta., and G. R. Galectins, Essentials of Glycobiology [Internet]. 4th edition, 10.1101/glycobiology.4e.36

17. Mkhikian, H., Sy, M., Dennis, J. W., and Demetriou, M. (2021) Galectins: The galectin-lattice: a decoder of bio-equivalent glycans. Glycoforum. 24, A10

18. Sato, S. (2023) Galectins: Why does galectin-3 have a unique intrinsically disordered region? “Raison d’être” for the disordered structure and liquid–liquid phase separation –Part 1–. Glycoforum. 26, A1

19. Sato, S. (2023) Galectins: Why does galectin-3 have a unique intrinsically disordered region? “Raison d’être” for the disordered structure and liquid–liquid phase separation –Part 2–. Glycoforum. 26, A8

20. Rabinovich, G. A. (2023) Galectins: The multifaceted roles of galectins: glycan-binding proteins with multiple personalities. Glycoforum. 26, A12

21. Troncoso, M. F., Elola, M. T., Blidner, A. G., Sarrias, L., Espelt, M. V., and Rabinovich, G. A. (2023) The universe of galectin-binding partners and their functions in health and disease. J. Biol. Chem. 299, 105400

22. Dennis, J. W. (2015) Many Light Touches Convey the Message. Trends Biochem Sci. 40, 673–686

23. Nabi, I. R., Shankar, J., and Dennis, J. W. (2015) The galectin lattice at a glance. Journal of cell science. 128, 2213–2219

24. Johannes, L., Jacob, R., and Leffler, H. (2018) Galectins at a glance. Journal of cell science. 131, jcs208884

25. Satoh, M. S., Rancourt, A., Boucher, E., St-Pierre, G., Hagiwara, K., Nakajima, K., Fillion, M., and Sato, S. (2024) N-Acetylglucosamine Facilitates Coordinated Myoblast Flow, Forming the Foundation for Efficient Myogenesis. bioRxiv

26. Cutler, A. A., Pawlikowski, B., Wheeler, J. R., Betta, N. D., Elston, T., O’Rourke, R., Jones, K., and Olwin, B. B. (2022) The regenerating skeletal muscle niche drives satellite cell return to quiescence. iScience. 25, 104444

27. Arecco, N., Clarke, C. J., Jones, F. K., Simpson, D. M., Mason, D., Beynon, R. J., and Pisconti, A. (2016) Elastase levels and activity are increased in dystrophic muscle and impair myoblast cell survival, proliferation and differentiation. Sci. Rep. 6, 24708

28. Haslett, J. N., Sanoudou, D., Kho, A. T., Bennett, R. R., Greenberg, S. A., Kohane, I. S., Beggs, A. H., and Kunkel, L. M. (2002) Gene expression comparison of biopsies from Duchenne muscular dystrophy (DMD) and normal skeletal muscle. Proceedings of the National Academy of Sciences of the United States of America. 99, 15000–15005

29. Ryczko, M. C., Pawling, J., Chen, R., Rahman, A. M. A., Yau, K., Copeland, J. K., Zhang, C., Surendra, A., Guttman, D. S., Figeys, D., and Dennis, J. W. (2016) Metabolic Reprogramming by Hexosamine Biosynthetic and Golgi N-Glycan Branching Pathways. Sci Rep. 6, 23043

30. Grigorian, A., Lee, S. U., Tian, W., Chen, I. J., Gao, G., Mendelsohn, R., Dennis, J. W., and Demetriou, M. (2007) Control of T Cell-mediated autoimmunity by metabolite flux to N-glycan biosynthesis. J. Biol. Chem. 282, 20027–20035

31. Grigorian, A., Araujo, L., Naidu, N. N., Place, D. J., Choudhury, B., and Demetriou, M. (2011) N-Acetylglucosamine Inhibits T-helper 1 (Th1)/T-helper 17 (Th17) Cell Responses and Treats Experimental Autoimmune Encephalomyelitis*. J. Biol. Chem. 286, 40133–40141

32. Takahashi, M., Inoue, K., Yoshida, M., Morikawa, T., Shibutani, M., and Nishikawa, A. (2009) Lack of chronic toxicity or carcinogenicity of dietary N-acetylglucosamine in F344 rats. Food and chemical toxicology : an international journal published for the British Industrial Biological Research Association. 47, 462–471

33. Lee, K. Y., Shibutani, M., Takagi, H., Arimura, T., Takigami, S., Uneyama, C., Kato, N., and Hirose, M. (2004) Subchronic toxicity study of dietary N-acetylglucosamine in F344 rats. Food and chemical toxicology : an international journal published for the British Industrial Biological Research Association. 42, 687–695

34. Reagan-Shaw, S., Nihal, M., and Ahmad, N. (2008) Dose translation from animal to human studies revisited. FASEB journal : official publication of the Federation of American Societies for Experimental Biology. 22, 659–661

35. Salvatore, S., Heuschkel, R., Tomlin, S., Davies, S. E., Edwards, S., Walker-Smith, J. A., French, I., and Murch, S. H. (2000) A pilot study of N-acetyl glucosamine, a nutritional substrate for glycosaminoglycan synthesis, in paediatric chronic inflammatory bowel disease. Aliment. Pharmacol. Ther. 14, 1567–1579

36. Sy, M., Newton, B. L., Pawling, J., Hayama, K. L., Cordon, A., Yu, Z., Kuhle, J., Dennis, J. W., Brandt, A. U., and Demetriou, M. (2023) N-acetylglucosamine inhibits inflammation and neurodegeneration markers in multiple sclerosis: a mechanistic trial. J. Neuroinflammation. 20, 209

37. McArdle, A., Edwards, R. H. T., and Jackson, M. J. (1994) Time course of changes in plasma membrane permeability in the dystrophin-deficient mdx mouse. Muscle Nerve. 17, 1378– 1384

38. Bulfield, G., Siller, W. G., Wight, P. A., and Moore, K. J. (1984) X chromosome-linked muscular dystrophy (mdx) in the mouse. Proc. Natl. Acad. Sci. 81, 1189–1192

39. Pernold, K., Iannello, F., Low, B. E., Rigamonti, M., Rosati, G., Scavizzi, F., Wang, J., Raspa, M., Wiles, M. V., and Ulfhake, B. (2019) Towards large scale automated cage monitoring – Diurnal rhythm and impact of interventions on in-cage activity of C57BL/6J mice recorded 24/7 with a non-disrupting capacitive-based technique. PLoS ONE. 14, e0211063

40. Iannello, F. (2019) Non-intrusive high throughput automated data collection from the home cage. Heliyon. 5, e01454

41. Szmyd, B., Rogut, M., Białasiewicz, P., and Gabryelska, A. (2021) The impact of glucocorticoids and statins on sleep quality. Sleep Med. Rev. 55, 101380

42. Sasai, K., Ikeda, Y., Fujii, T., Tsuda, T., and Taniguchi, N. (2002) UDP-GlcNAc concentration is an important factor in the biosynthesis of beta1,6-branched oligosaccharides: regulation based on the kinetic properties of N-acetylglucosaminyltransferase V. Glycobiology. 12, 119–127

43. Rahman, A. M. A., Ryczko, M., Nakano, M., Pawling, J., Rodrigues, T., Johswich, A., Taniguchi, N., and Dennis, J. W. (2015) Golgi N-glycan branching N-acetylglucosaminyltransferases I, V and VI promote nutrient uptake and metabolism. Glycobiology. 25, 225–240

44. Gagneux, P., Hennet, T., and Varki, A. (2022) Biological Functions of Glycans. 10.1101/glycobiology.4e.7

45. Collins, B. C., Shapiro, J. B., Scheib, M. M., Musci, R. V., Verma, M., and Kardon, G. (2024) Three-dimensional imaging studies in mice identify cellular dynamics of skeletal muscle regeneration. Dev. Cell. 59, 1457–1474.e5

46. Webster, M. T., Manor, U., Lippincott-Schwartz, J., and Fan, C.-M. (2016) Intravital Imaging Reveals Ghost Fibers as Architectural Units Guiding Myogenic Progenitors during Regeneration. Cell Stem Cell. 18, 243–252

47. Wooddell, C. I., Radley-Crabb, H. G., Griffin, J. B., and Zhang, G. (2011) Myofiber Damage Evaluation by Evans Blue Dye Injection. Curr. Protoc. Mouse Biol. 1, 463–488

48. Sato, S., Rancourt, A., and Satoh, M. S. (2024) Cell Fate Simulation Reveals Cancer Cell Features in the Tumor Microenvironment. J. Biol. Chem. 10.1016/j.jbc.2024.107697

49. Tsonaka, R., Signorelli, M., Sabir, E., Seyer, A., Hettne, K., Aartsma-Rus, A., and Spitali, P. (2020) Longitudinal metabolomic analysis of plasma enables modeling disease progression in Duchenne muscular dystrophy mouse models. Hum. Mol. Genet. 29, 745–755

