## Supplementary figures and images for "Improvement of Spontaneous Locomotor Activity in a Murine Model of Duchenne Muscular Dystrophy by N-Acetylglucosamine Alone and in Combination with Prednisolone"

### Supplemental figure

#### Sup.Fig. 1

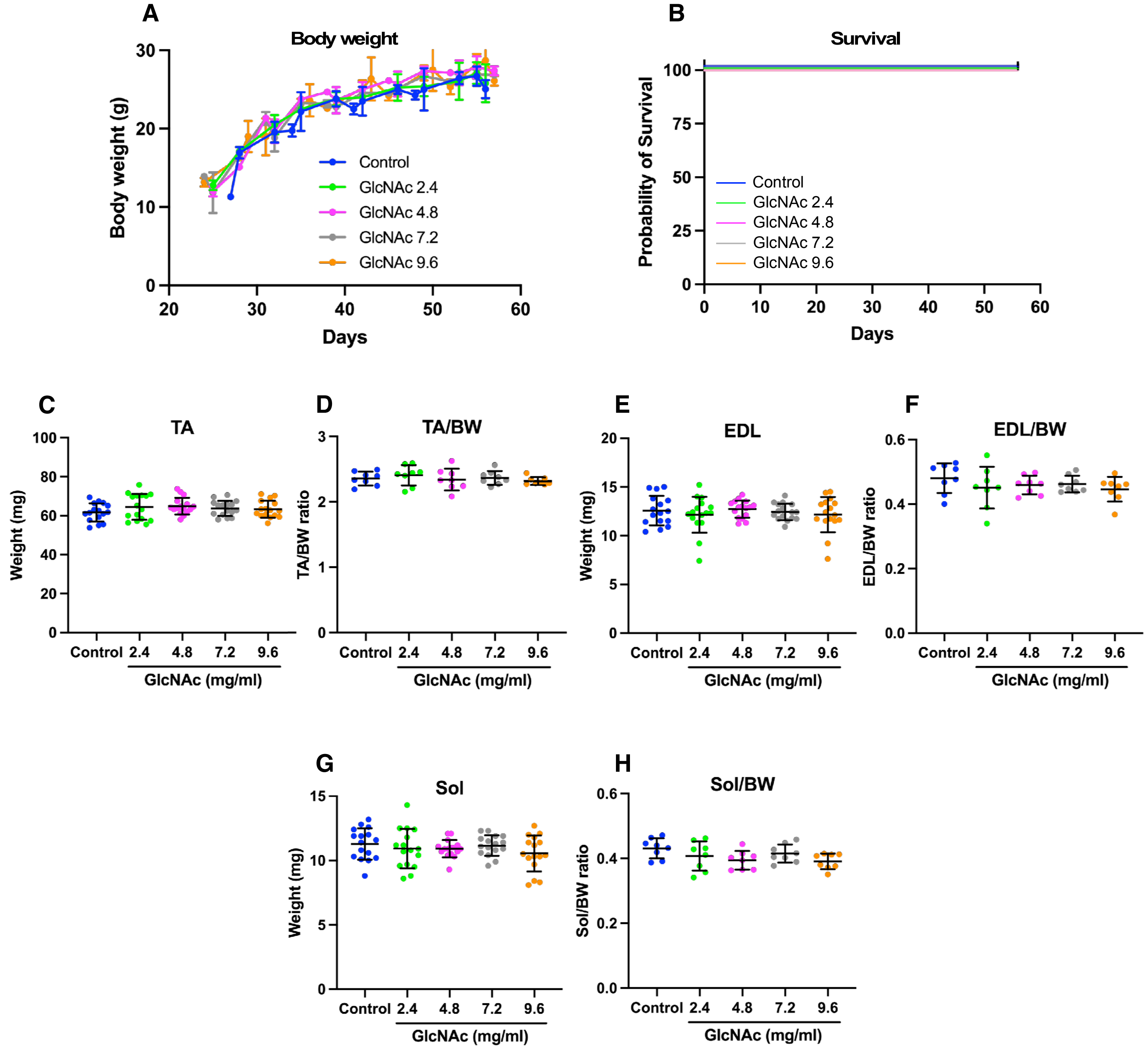


Sato et al.

#### Sup. Fig. 2

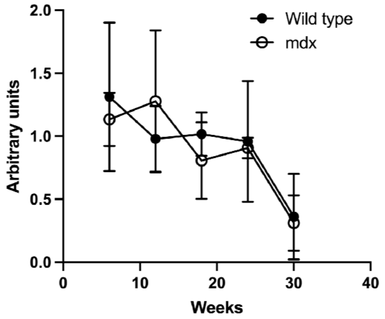


Sato et al.
